# Improving 3D Edge Detection for Visual Inspection of MRI Coregistration and Alignment

**DOI:** 10.1101/2022.09.14.507937

**Authors:** Chris Rorden, Roger Newman-Norlund, Chris Drake, Daniel R. Glen, Julius Fridriksson, Taylor Hanayik, Paul A. Taylor

## Abstract

Detecting and visualizing edges is important in several neuroimaging and medical imaging applications. For example, it is common to use edge maps to ensure the automatic alignment of low-resolution functional MRI images to match a high-resolution structural image has been successful. Specifically, software toolboxes like FSL and AFNI generate volumetric edge maps that can be particularly useful for visually assessing the alignment of datasets, overlaying the edge map of one on the other. Therefore, edge maps play a crucial role in quality assurance. Popular methods for computing edges are based on either the first derivative of the image as in FSL, or a variation of the Canny Edge detection method as implemented in AFNI. The crucial algorithmic parameter for adjustment for each of these methods relates to the image intensity. However, image intensity is relative and can be quite variable in most neuroimaging modalities. Further, the existing approaches do not necessarily generate a closed edge/surface, which can reduce the ability to determine the correspondence between a represented edge and another image. We suggest that using the second derivative (difference of Gaussian, or DoG) of the image to generate edges resolves both these issues. This method primarily operates by specifying a spatial scale of interest (which is typically known in medical imaging) rather than a contrast scale, and creates closed surfaces by definition. We describe some convenient implementation features (for both efficiency and visual quality) developed here, and we provide open source implementations of this method as both online and high performance portable code. Finally, we include this method as part of both the AFNI and FSL software packages.

## Introduction

Edge detection identifies key features in an image, which, unless otherwise stated, means a 3D volume here and below. Popular neuroimaging toolboxes such as AFNI (Cox, 1996) and FSL (Smith et al., 2004) typically include bespoke tools for edge detection, where they are commonly used to generate an outline of an image or of specific features within the image. For example, locations of contrast showing tissue boundary delineation. Additionally, these edge maps can be overlaid or combined with another image to help the user determine the relative alignment of anatomical structures within pairs of images. Edge detection is a common component of image processing, and numerous algorithms have been proposed in the literature to address certain challenges in a given application (e.g., low contrast images, noisy images, etc.). Here we discuss the properties of several popular edge detection methods and suggest that the second derivative is well suited for many medical imaging applications.

In the field of neuroimaging, a common application for edge maps is to evaluate image registration, where one volumetric image is aligned to another. This can occur between separate acquisitions of the same subject (often containing different contrast mechanisms and/or nonlinear distortion patterns), as well as between different subjects or between a subject and a standard template. These automated image registration methods iteratively adjust the position of an input (source) image to match the target (base) image, attempting to minimize a cost function, which estimates the alignment. Different cost functions have been suggested and evaluated for quality and robustness (for example, in MRI, see Saad et al., 2009). However, all algorithms risk becoming trapped in a local minimum where the final alignment is suboptimal or entirely incorrect (Brett et al., 2001), particularly when the images contain noise, low contrast or poor initial alignment. Moreover, alignment results by definition cannot be quantitatively assessed independently of the cost function: if a separate quantity *could* provide such an evaluation of the goodness-of-alignment, then it should be used or incorporated into the cost function itself. Therefore, visual inspection of automated alignment algorithms is an important step of registration and quality control (Saad, 2009). At the same time, it can be challenging to see details throughout both images simultaneously. One or both images can be inherently obscured due to large noise, acquisition artifacts, superimposed brightness patterns, poor image contrast, and differing image contrasts (e.g., bright ventricles in the source image, but dark ventricles in the base image). Overlaying edge maps from the input images on top of the target image (or vice versa) can provide efficient and useful visual information to researchers in quality assessment reports. Alignment and edge maps used in this way can also promote detection of other forms of data conversion and processing errors, such as the relative left-right flipping of images (e.g., Glen et al., 2020).

At present, the quality assessment images generated by some popular methods can be challenging to interpret, especially with respect to differentiating anatomical contours when the edge algorithm creates disjointed boundaries. As noted above, visual inspection is crucial. Automated methods do not always perform as intended (Raslau et a., 2018), and reliance on automated methods for quality assurance is problematic. Indeed, a grossly distorted solution caused by the presence of a lesion (Brett et al., 2001) or susceptibility artifacts (Raslau et al., 2018) can often have better alignment with these cost functions than solutions that are actually more accurate. Therefore, having reliable edge detection and display algorithms is essential for neuroimaging interpretation and processing.

Creating edge maps can be particularly challenging for MRI modalities that are not optimized to maximize intensity contrast between different tissue types. The parameters of a structural T1-weighted scan are typically tuned to highlight contrast between gray matter (GM), white matter (WM) and CSF, and the dataset is typically acquired with high spatial precision (relative to the size of easily identified anatomical structures). In contrast, modalities such as functional magnetic resonance imaging (FMRI), diffusion tensor imaging (DTI) and arterial spin labeling (ASL) are designed to derive properties over time or multiple volumes (such as brain function, white matter bundle orientation, blood flow, respectively). The acquired images often exhibit low signal-to-noise (SNR), poor tissue contrast, poor intensity homogeneity, signal dropout and low spatial resolution, all of which pose unique challenges to edge-detection algorithms as will be discussed below. To improve alignment, previous efforts have altered the imaging acquisition protocol to enhance spatial contrast explicitly (e.g., Gonzalez-Castillo et al., 2013).

In this work we briefly describe three seminal edge detection methods: first derivative gradients, second derivative zero-crossing (which we refer to as Difference of a Gaussian, DoG) and the multi-stage Canny algorithm. We describe how popular neuroimaging tools have previously identified edges in 3D images, with FSL using the first derivative and AFNI having adapted a portion of the Canny algorithm. We then propose that the DoG may be particularly well suited to this application, with advantages over both other methods. First, the DoG generates closed loops, which can help identify the shape of 3D anatomical structures when observing 2D slices. Second, the DoG is rapid and easy to implement for 3D images. This contrasts, for example, to the latter stages of the full Canny filter (Bähnisch et al., 2009). Finally, all algorithms have some parameter dependence, but the primary setting for the DoG is based on the spatial size of features of interest, which is usually known for each imaging application. In contrast, the other methods typically require intensity-based thresholds, which vary by scanner, across volume and are generally not well-calibrated for most MRI modalities. We describe our DoG implementation(s), including new optimization steps. We provide examples for various popular imaging modalities, illustrating that the DoG provides a robust and easily interpretable solution. As developers for and in conjunction with several software packages, we have now implemented DoG in AFNI, FSL, Matlab/SPM, Python/nibabel, niimath and a new web-based viewing tool called NiiVue.

### Background: Evolution of Edge Detection Operators

Historically, several computational methods have been devised for edge detection, originally for 2D images and then generalizing to 3D. We briefly describe three generations of methods. Edges typically correspond with image gradients: locations where the brightness or color in a pixel or voxel differs from its neighbors. Most neuroimaging modalities generate scalar grayscale images where brightness is explicit. For multi-component color images, (e.g. where each voxel is defined by Red, Green and Blue values) one can choose a relative luminance transform to define a scalar brightness. We describe some methods below, occasionally with 2D-based examples for simplicity but each of them has been generalized to 3D.

The 2D Roberts cross operator (1965) is a pioneering edge-detection solution based on the first derivative of the image’s intensity values. This operator defines the strength of a horizontal gradient *Gx* as the difference between a pixel’s intensity and the intensity of a neighbor that shares its lower right corner; the 2×2 kernel representation for this is [1 0; 0 −1], using the Matlab convention of describing numerical matrices with a semicolon as the row delimiter. The vertical gradient *Gy* is the difference between a pixel’s intensity and that of its lower left neighbor, with kernel [0 1; −1 0]. Note that due to the kernel structure the gradient values appear on one of the pixels used in the calculations, rather than between them, effectively shifting the gradient by half a pixel along each dimension. That is, the edge itself is an image, and so it typically exists on one or more of the pixels used in the kernel of the gradient calculation. To remove this spatial bias, one could alternatively sample neighbors to either side of the pixel of interest, using a central tendency operator [1 0 −1] to estimate *Gx* by comparing the pixels directly to the left and right of the pixel of interest, and [1; 0; −1] to estimate *Gy* by comparing the neighbors directly above and below. With either operator, the overall gradient magnitude G is given by sqrt ((*Gx*)^2 + (*Gy*)^2), and the resulting gradient image can be used directly or thresholded in some form to create edges. However, these operators are particularly sensitive to noise (Gonzalez and Woods, 1992). The Prewitt (1970) and Sobel (1970) operators expand the kernels to examine neighbors sharing either faces or corners. These reformulations provide some smoothing to reduce noise susceptibility at the cost of some spatial sensitivity. Typically, Sobel is preferable to Prewitt, as the former is more effective at localizing edges and attenuating aliasing at essentially identical computational cost (Spontón and Cardelino, 2015).

Until the late-1970s and early-1980s, most edge detection algorithms relied on small-scale operators that recursively examined pixel variations between neighbors or within relatively small neighborhoods, with no consideration for edge characteristics and noise content (see Gonzalez and Woods, 1992). In 1980, Marr and Hildreth (1980) introduced a seminal edge-finding algorithm which was based on the second derivative of image intensity and built on earlier work of Wilson and Giese (1977). This algorithm introduced the Laplacian of a Gaussian (LoG) function, which contains parameters that can be tuned to select features at a given scale sigma, thereby attenuating local noise. Specifically, the function blurs the image twice with separate Gaussians of scale sigma and slightly larger sigma+epsilon, producing a “Mexican hat operator” that attenuates features smaller than sigma, and which is robust against ringing artifacts (Gonzalez and Woods, 1992). Using this technique, edges from the original volume can be identified based on the zero crossings of the convolved image. Note that all the produced zero-crossings by definition form closed contours, analogous to contour lines on an elevation map: the edge is the transition between bright islands (blobs) and dark regions in the convolved image.

While we later argue that these closed contours are typically a positive attribute for neuroimaging applications, we should also note that in some other applications the “spaghetti effect” generated by this technique can be counterproductive (Gonzalez and Woods, 1992). We emphasize that edge techniques have strengths and weaknesses that should be judged for each application. In the present application, avoiding the “spaghetti effect” effectively places a boundary on the parameter settings for an application, which is generally straightforward to avoid. Additionally, Marr and Hildreth (1980) noted that the LoG itself is difficult to compute. However, there is a close approximation that can be computed rapidly, using the Difference of a Gaussian (DoG), and this is typically what is adopted in implementations. Therefore, for brevity, we refer to the Marr and Hildreth method with edges based on second derivative zero-crossing as “DoG”.

In many applications, the Canny edge detector (1986) was considered to be superior to prior methods, including the Marr and Hildreth method, and the Canny method quickly became a tool of choice for applications relying heavily on accurate edge detection (Gonzalez and Woods, 1992). This algorithm uses a five-stage approach to identify edges. First, the image is blurred to attenuate high frequency noise (typically using a Gaussian blur). Next, first derivative gradients of this new image are calculated, quantifying how rapidly image intensity varies between pixels, traditionally using a Prewitt (1970) or Sobel (1970) filter. Next, non-maximum suppression is used to identify and preserve local maxima. Fourth, thresholds are applied to identify edges. Here two thresholds are applied: voxels with extremely strong gradients are identified as edges, while those with moderately strong thresholds become candidate locations that may be classified as edges in the subsequent stage. Finally, one conducts edge tracking by hysteresis, retaining moderate edges that improve the continuity of stronger ones and removing those that do not. The Canny method does not require edges to form closed loops. Note how this method relies on several steps, each typically having multiple parameters of scale, contrast or magnitude; thus, while it can be tuned to high accuracy for a given family of inputs, it can be challenging to retain high quality across a wide range of differing images.

### First-Derivative Method (used in FSL)

When generating coregistration quality assurance images, FSL identifies edges based on the first derivative of image intensity (via “slicer”). The outline images created by FSL are computed on 2D images sampled from a 3D volume (e.g. sagittal, coronal and axial images are each computed in plane). Specifically, FSL estimates a voxel gradient using the 2D gradient (central tendency) method previously described. Subsequently, the 2nd and 98th percentile of all the gradients in the image are computed, and the gradient map is converted into a binary edge map using the mean of these two percentiles as a threshold. This formula does not necessarily generate closed loops, and it will mask weak gradients that are present in many images (e.g. low contrast between tissues inside the brain relative to the strong gradients generated at the boundary between the brain and empty space which are created during FSL’s brain extraction).

### Canny Method (used in AFNI)

AFNI’s 3dedge3 program adapts the Canny edge detection algorithm, which was originally described for 2D images, to 3D using recursive filters (Deriche, 1987; Monga et al., 1991). It should be noted that the hysteresis stage of the Canny algorithm is especially complicated when applied in three dimensions (Bähnisch et al., 2009), and therefore popular 3D implementations do not typically use this step, including AFNI’s 3dedge3. A specific challenge faced when applying hysteresis thresholding to medical images is the issue of the varying contrast for edges, which makes automatically selecting the two thresholds in the Canny algorithm challenging. However, the edges estimated by 3dedge3 have still been useful for alignment purposes and are efficiently calculated, though perhaps containing more ‘noisy’ edges than the full algorithm would produce. We also note that, rather than generating a binary edge, 3dedge3 outputs a volume whose edge map has continuous intensity representing the magnitude (i.e. strength) of the edge, though it can also be displayed as a binary image (e.g. Saad et al., 2009). This approach makes it possible for users to manually adjust the gradient strength or manually threshold the output gradient image to highlight the most well-defined edges, if desired, in place of the hysteresis editing. However, again edges are not guaranteed to be closed loops, and the algorithm can be sensitive to underlying brightness patterns.

### DoG: Second Derivative Zero-Crossing Method (proposed)

We suggest that the DoG (second derivative, zero-crossing) method described by Wilson and Giese (1977) and Marr and Hildreth (1980) seems particularly well suited to the task of creating coregistration quality assurance images used by MRI researchers. Both the first derivative and Canny methods require threshold(s) based on gradient intensity. However, different tissue boundaries may have different relative contrasts, so a single threshold may be hard to define (as illustrated in the Figures described below). Likewise, due to effects like image inhomogeneity the relative tissue intensities can gradually change across the image, requiring the application of different thresholds at different image locations. Fundamentally, image intensity in most MRI sequences has relative arbitrary units (unlike CT, where tissue intensity is typically in known Hounsfield units, which have physical relevance), making it difficult to tune brightness-dependent parameters, outside of using mean intensity (which can be more or less influenced by field of view size and other image features).

The primary adjustable parameter of the Marr and Hildreth (1980) algorithm is the ‘inner Gaussian’ sigma, which can be set according to the spatial size of the focal features within an image. This is a relatively convenient parameter to be able to set, particularly for visually assessing biological/anatomical alignment, since the minimal scale size of important features is typically known. For example, in human imaging the thickness of the gray matter that comprises the human cortex (and forms the *de facto* outline of the brain in two-dimensional axial, sagittal and coronal slices) is approximately 2-2.5mm (Fischl and Dale, 2000). Pragmatically, in adult human neuroimaging features smaller than 2mm are unlikely to be of interest for coregistration, providing a useful “default” threshold for this algorithm in human neuroimaging to focus on GM while also suppressing extraneous, subscale features. When imaging of other species, an analogous scale size is similarly known. Additionally, users can also explicitly set this parameter to an appropriate length scale for any applications (e.g., high resolution subcortical scans, ultra-high field imaging that discriminates cortical layers, animal scans of smaller brains, etc.). In the Discussion, we note how image voxel dimensions often provide a useful proxy of spatial scale of interest for anatomical images, so a reasonable default for general, cross-species applications is possible.

Further, the closed contours formed by second derivative zero-crossing algorithms are ideally suited for matching the outlines of anatomical structures, especially when viewed slicewise. Specifically, when a user examines the edges of one image on top of the other, the shape of the edges generated by this operator can help identify what structure the edge refers to.

## Methods

### DoG algorithm and implementations

Here we briefly discuss the original edge algorithm of Wilson and Giese (1977) and Marr and Hildreth (1980), which is shown schematically in Figure 1. Additionally, we discuss modern updates to features, which have computational and/or interpretational benefits. We note parameters within the algorithm, describing the effects of each and how defaults can be chosen. We have implemented this algorithm in several tools (AFNI, FSL, Matlab, niimath, Python). There are slight differences among these implementations (defaults, optionality, etc.), and we note these when relevant.

**Figure 1.**
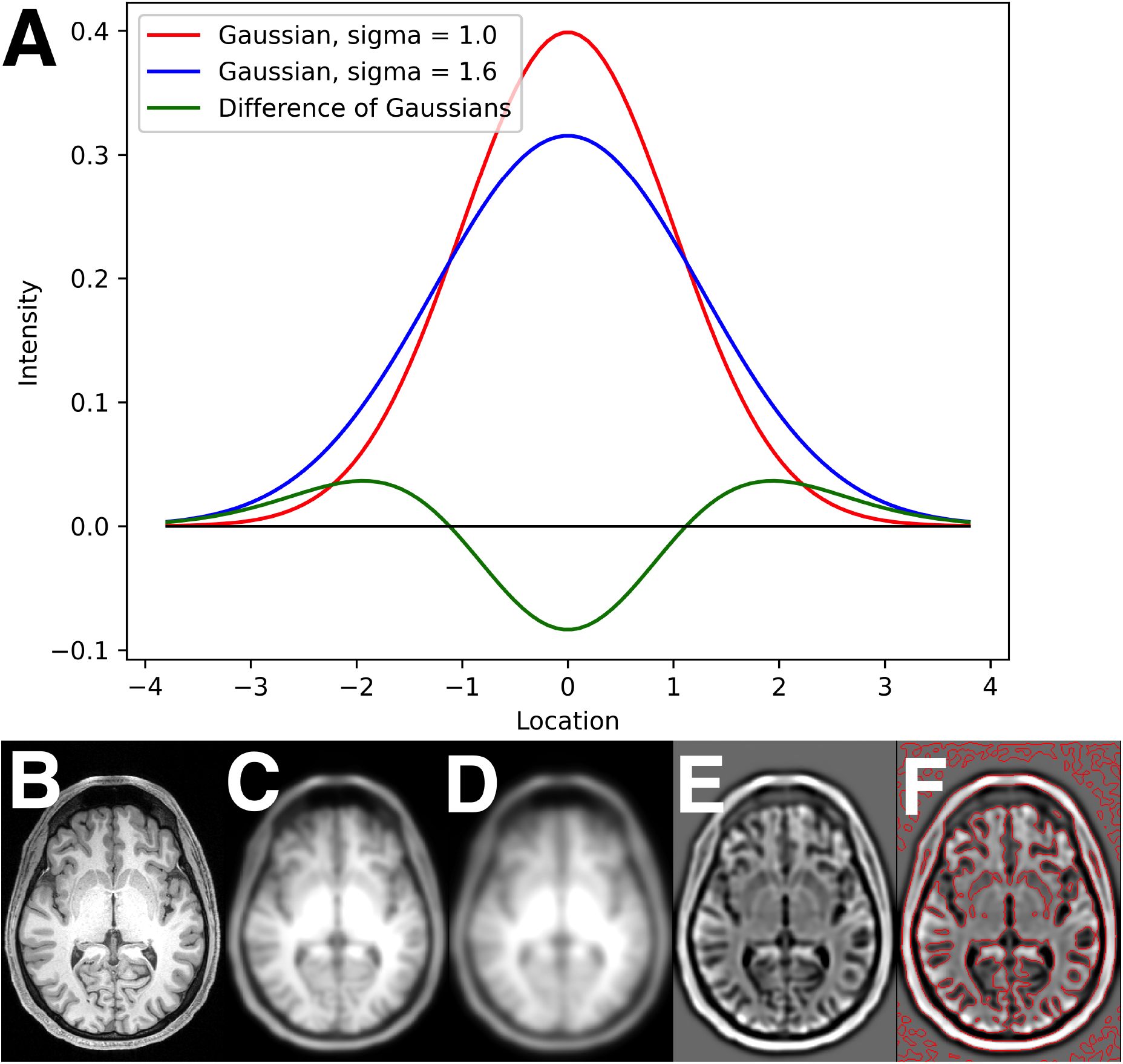
Illustration of the Difference of Gaussian edge detection. A) A schematic plot showing the narrower “inner” (σ = 1.0, red) and wider “outer” (σ = 1.6, blue) Gaussian kernel; the horizontal axis ticks are in units of σ, and the vertical axis shows intensity amplitude. The difference between these two kernels is shown in green. B) Initial T1-weighted MRI image. C) Image blurred with sigma_inner = 2.0. D) Image blurred with sigma_outer = 3.2mm Gaussian. E) Difference between images, C -D. F) Zero-crossings of image E superimposed in red.

In attempting to identify particularly beneficial edge-detection algorithms for 3D image alignment/coregistration and quality control/visualization, we initially examined the LoG- or DoG-based approaches of Wilson and Giese (1977) and Marr and Hildreth (1980). The LoG function is not separable, and therefore it is computationally expensive to apply to 3D images, as the number of underlying calculations increases with N^3^. Even with modern computing this is undesirable, and numerous strategies to accelerate this filter have been described (e.g., Chen et al., 1987). However, there are two popular, simple, and computationally efficient methods that can *approximate* this function well, even for 3D images. The first method involves blurring the data with a Gaussian lowpass filter (which is a spatially separable procedure) and subsequently applying a Laplacian with a 3×3×3 kernel to the image. The second method, which is adopted in the programs implemented here, involves using the Difference of Gaussians (DoG) method, which had been previously described by Wilson and Giese (1977) in the context of psychophysical studies. Here one applies a pair of “inner” and “outer” Gaussian blurs (which is highly efficient, due to separability and Gaussian additive properties) to the source image (see Figure 1A,C,D), with the result being the difference between these two images (Figure 1E). The value of the inner sigma (or “sigma_rad”) is the primary parameter to set in this algorithm, as above noted, which can be set based on the spatial scale of interest. It is important to note that the scale width of the inner Gaussian is two times the inner radius value; therefore, 2*sigma_rad is the appropriate quantity to make less than a minimal structural length.

The ratio of these two Gaussians (the “ratio_sigma”) is a secondary parameter that affects the amount of detail traced with edges. Wilson and Giese (1977) suggested a value of ratio_sigma=1.5, and Marr and Hildreth (1980) suggested 1.6 in their work. Neither of the initially suggested values of ratio_sigma were definitively derived, and it is possible that this parameter could be application-dependent. It must be noted, though, that the similarity of DoG and LoG functions *decreases* as this ratio increases, so one would not expect a very large value---perhaps even values of 1.5-1.6 might be considered surprisingly large. Therefore, we investigated its effect here.

Having identified a quick and robust method for implementing DoG, the next step in the edge algorithm is to define locations of zero-crossings in the DoG image, which become the edges themselves (Figure 1F). There are several approaches for computing edge crossings, including those that offer sub-pixel accuracy (Huertas & Medioni 1986). One conceptually simple approach is to count a voxel as being an edge if the location is positive in the DoG and yet shares a neighbor with a negative DoG value (or vice versa, with a negative pixel neighboring a positive one). This would preferentially select the edge from the “bright” (or “dark”, respectively) region in the original image, which in some applications might be a usefully consistent choice. A related alternative that would not select one side consistently would be to define a zero crossing edge as a voxel that shares a neighbor with both opposite polarity and has greater magnitude/intensity. This is the strategy adopted in our niimath C program implementation. This method is fast in low level languages, but the reliance on conditionals and non-vectorized loops makes it difficult to implement efficiently in most higher-level languages. On the other hand, our AFNI, FSL, Matlab, and Python (3dedgedog, also in C) programs use a three step approach to identify zero crossings. First, binarize the DoG image, so that voxels >=0 become 1 and the rest are zero. Then, perform the Euclidean Distance Transform (EDT) on this image, which calculates each voxel’s distance from the mask boundary; we utilize the highly efficient algorithm by Felzenszwalb and Huttenlocher (2012) for this step. Finally, select voxels with a distance of one edge length (or some other chosen value) to define the boundary. This basic method provides consistent boundaries (e.g., all on one side of the contrast edge) of uniform thickness (which adapts to anisotropic voxels and, if needed, can be adjusted), while maintaining computational efficiency.

The above steps define the central DoG algorithm. We note a few options and variations that are also implemented in specific programs, for various use cases. The main algorithm finds edges defined in the 3D volume, but sometimes it can be useful to find only edges within a 2D plane (e.g., for slicewise viewing of a volume); the above algorithm translates directly to this, and both the niimath and AFNI implementations have options for calculating slicewise edges through any of the main image orientations. Additionally, the input maps can be “automasked”, to avoid showing edges of background/noise, which can be visually distracting. AFNI’s 3dedgedog has options for selecting a particular side of a boundary: positive (=higher contrast side), negative (=lower contrast side), both (=positive+negative bounds), and both-signed (= same as both, but higher and lower contrast sides have positive and negative values, respectively). 3dedgedog can also display an approximate magnitude of each boundary location, estimated as the standard deviation in that voxel’s immediate neighborhood. Finally, 3dedgedog allows the user to select values of ratio_sigma and sigma_rad, as well as to define the kernels anisotropically or in terms of the number of voxels.

## Testing data

To demonstrate the properties of both existing and proposed edge detection strategies, we utilize the following publicly available MRI datasets. Firstly, we use the standard T1-weighted (T1w) anatomical dataset from the AFNI Bootcamp^1^, referring to this dataset as “sub-001”. This was acquired at 3T with standard resolution of near 1mm resolution (voxel = 1 × 0.938 × 0.938 mm^3^); this is particularly useful for examining DoG algorithm edges with changing parameters.

Additionally, we used a dataset containing multiple modalities from a single individual (“sub-002”), being representative of typical T1w and T2*-weighted images that would be acquired as part of a standard research neuroimaging protocol: T1w, arterial-spin labeling (ASL), diffusion weighted imaging (DWI) and functional MRI (fMRI). All data were acquired on a 3T Siemens Prisma Fit Scanner. The fMRI scan used a 90×90 matrix, 50 slices, 2.4mm isotropic voxels, TR=1650ms. The DWI sequence used a 140×140 matrix, 80 slices, 1.5mm isotropic voxels, TR=5700ms. The ASL (arterial-spin labeling) scan used a 70×70 matrix, 14 slices, 3.0×3.0×7.5mm voxels, TR=2500ms. The T1w sequence was acquired at high resolution with 0.8mm isotropic voxels.

We used AFNI’s 3dAllineate to rigidly align the first volume of each EPI dataset to the T1w volume, resampling to the latter’s grid. No corrections for EPI artifacts were applied, as our goal was to generate QA images that show regions with both good and poor edge alignment.

## Results

### DoG parameter investigation

We calculated DoG-derived edges for the standard T1w anatomical (approx. 1mm isotropic voxels) of sub-001, investigating part of the parameter space of the algorithm. The sigma_rad (= inner Gaussian sigma) took values from 0.6-2.6 mm, and ratio_sigma (= ratio of outer to inner Gaussian sigmas) took values from 1.2-2.0, using 3dedgedog. The results of these calculations are shown in Figure 2, for a zoomed-in portion of a sagittal slice that contains several scales of features and image contrast.

**Figure 2.**
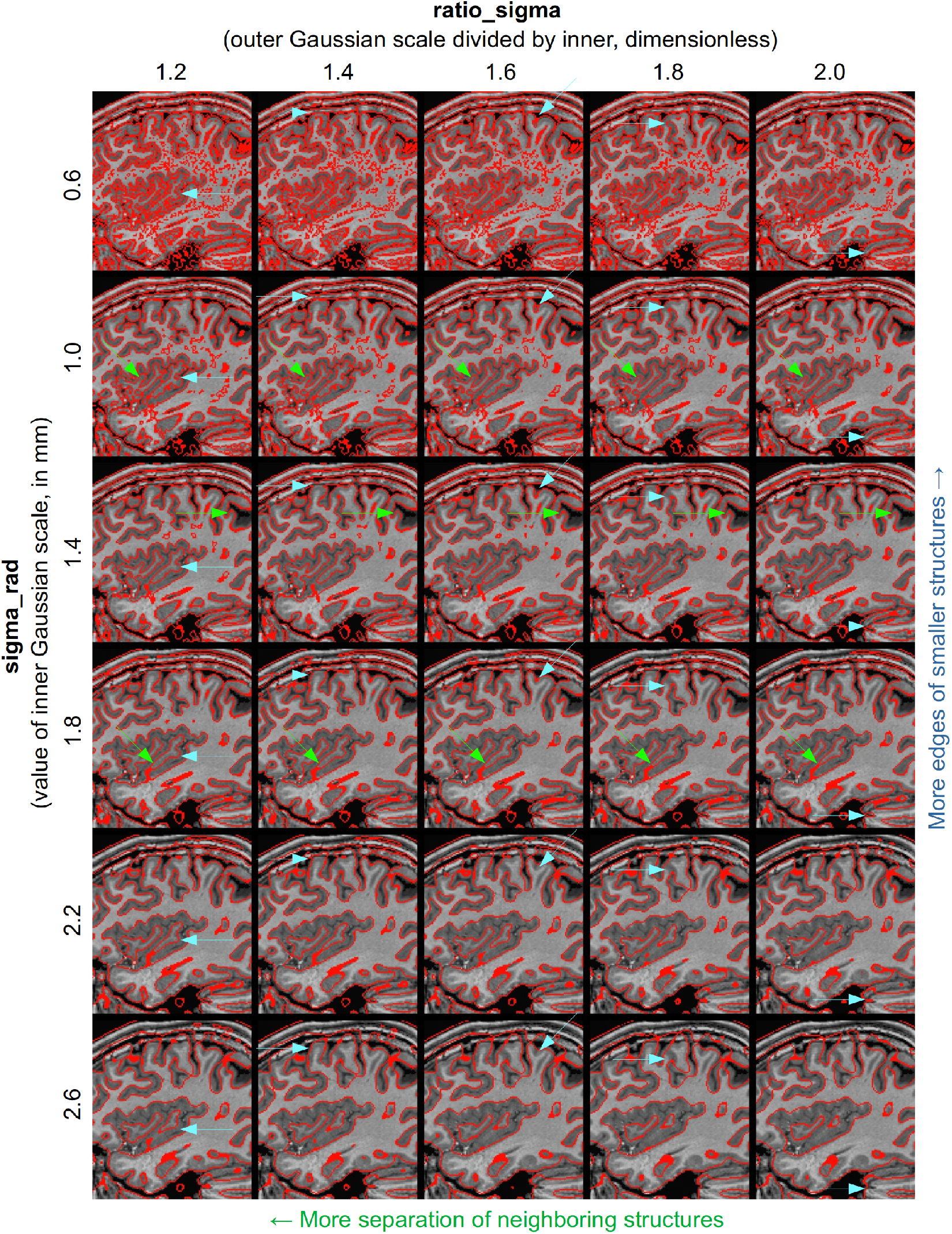
A grid of 3dedgedog results showing the effects of two main parameter choices in the DoG algorithm: the sigma_rad (radius of inner Gaussian) and the ratio_sigma (ratio of outer to inner Gaussian sigmas). The underlaid dataset, which was the input, is a standard T1w anatomical (sagittal slice at *x* = 40R). Blue arrows highlight example features where results differ noticeably with sigma_rad: edges of finer features can be observed as this parameter decreases. This parameter should be smaller than the desired scale of structure to observe (here, <2-2.5mm, which is the typical GM thickness), though as the value gets smaller one might consider some of the observed features to be effectively noise. Green arrows highlight example features where results changed noticeably with ratio_sigma: smaller and finer features can “melt away” as this parameter increases. This likely reflects the fact that as ratio_sigma approaches unity, the DoG method better approximates the LoG. Interestingly, it is often difficult for the DoG approach to show both the gray-white boundary and the pial boundary (“outer” GM bound), with the latter typically observed with other large-contrast boundaries.

The effects of altering sigma_rad are observed by comparing results in a given column. As expected, edges of smaller features are visible as sigma_rad decreases; as sigma_rad increases, more boundaries are either missed or incorrectly merged across unrelated structures. Blue arrows highlight representative examples of features changing with this parameter. It should also be noted that fine, low-contrast features also appear at the low sigma_rad values, such as within-WM edges; these might be considered noisy, for judging alignment, and perhaps could be filtered with considerations of non-binarized boundary magnitude.

The effects of altering ratio_sigma can be observed across a row. These effects tend to be subtle, but there are notable changes with this parameter. As ratio_sigma increases, finer features tend to decrease in size, notably at sharp curves or finer structures. Green arrows highlight representative examples of this.

Overall, for smaller sigma_rad and ratio_sigma, the edges appear to highlight many of the important edge features within the image. For assessing visual alignment of brains, one typically focuses on locations of tissue contrasts, such as the gray-white boundary and pial surfaces, and ventricle and other CSF boundaries. Sulcal and gyral structures are clearly visible here, though at these scales the edges typically show only one of the GM boundaries. For alignment purposes, the parameters used for the upper left panels of Figure 2 should provide the best representations (low ratio-sigma and low sigma_rad).

### Comparison of edges across software and modalities

The results of estimating edges for different MRI modalities (which have very different tissue and image contrast) with various algorithms used in different software toolboxes for sub-002 are shown in Figures 3-5 (for EPI-derived fMRI, DWI and ASL acquisitions, respectively, each with T1w). In each figure, the left set of panels shows the EPI-derived sequence as underlay with the T1w edges overlaid, and the right set of panels shows the T1w data underlaid and the EPI-derived edges overlaid; the top row shows FSL’s first derivative method (via “slicer”), the middle row shows AFNI’s Canny method (via “3dedge3”), and the bottom row shows AFNI’s new DoG method (our different implementations create nearly identical results).

**Figure 3.**
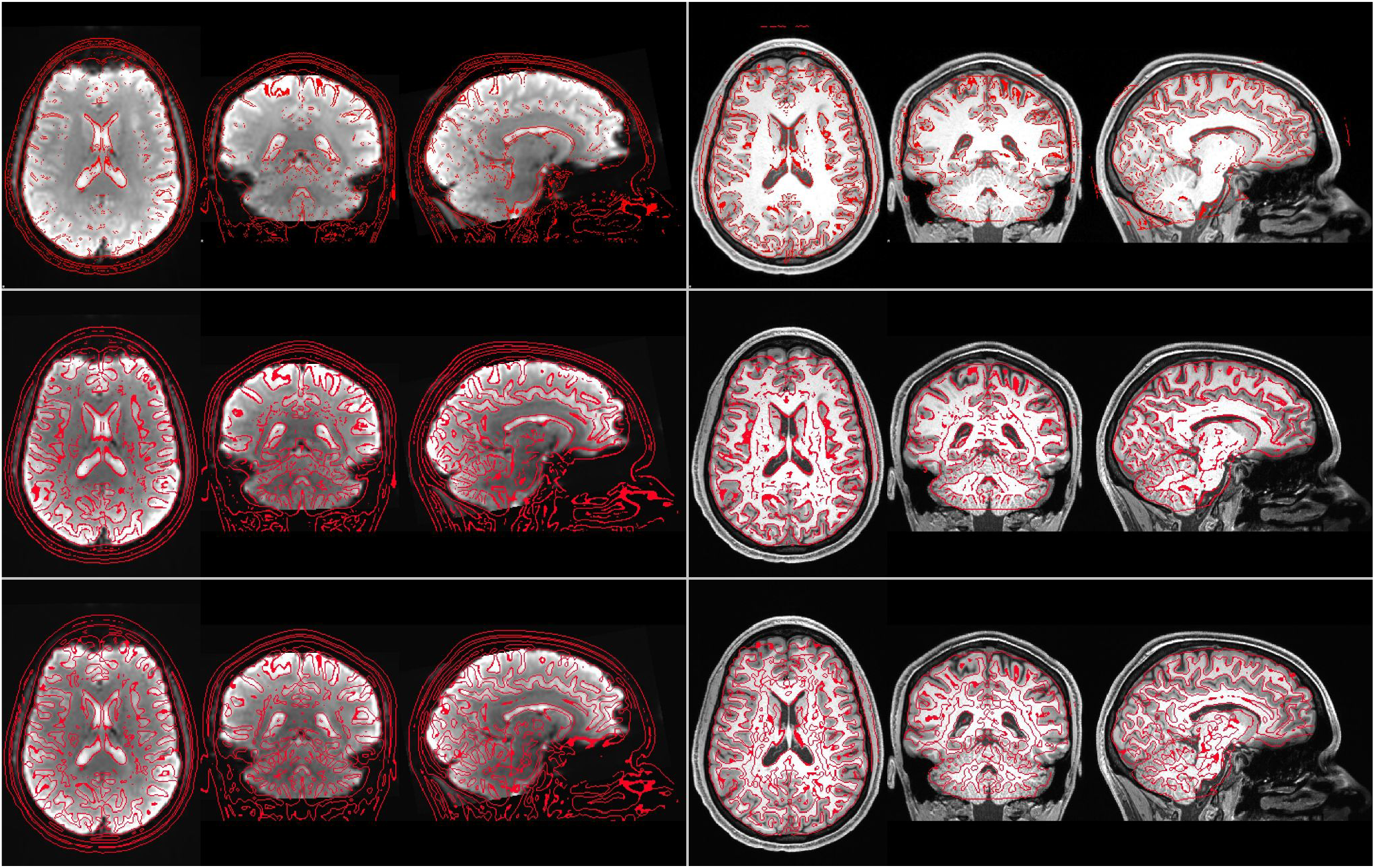
Example of rigid-body coregistration of a subject’s fMRI EPI to a T1w volume using different edge approaches (from top row, downward): FSL’s slicer (first derivative-based method), AFNI’s 3dedge3 (Canny-based method) and AFNI’s 3dedgedog (new DoG-based method, with sigma_rad=1.4 and ratio_sigma = 1.4). In the left panels, the EPI-based volume is underlaid beneath edges from the T1w volume, and in the right panels, the T1w is beneath EPI-derived edges. The sagittal, coronal and axial images are centered at (−7.9, 42.4, 20.8) in RAI-DICOM coordinates, and each image left = subject right.

**Figure 4.**
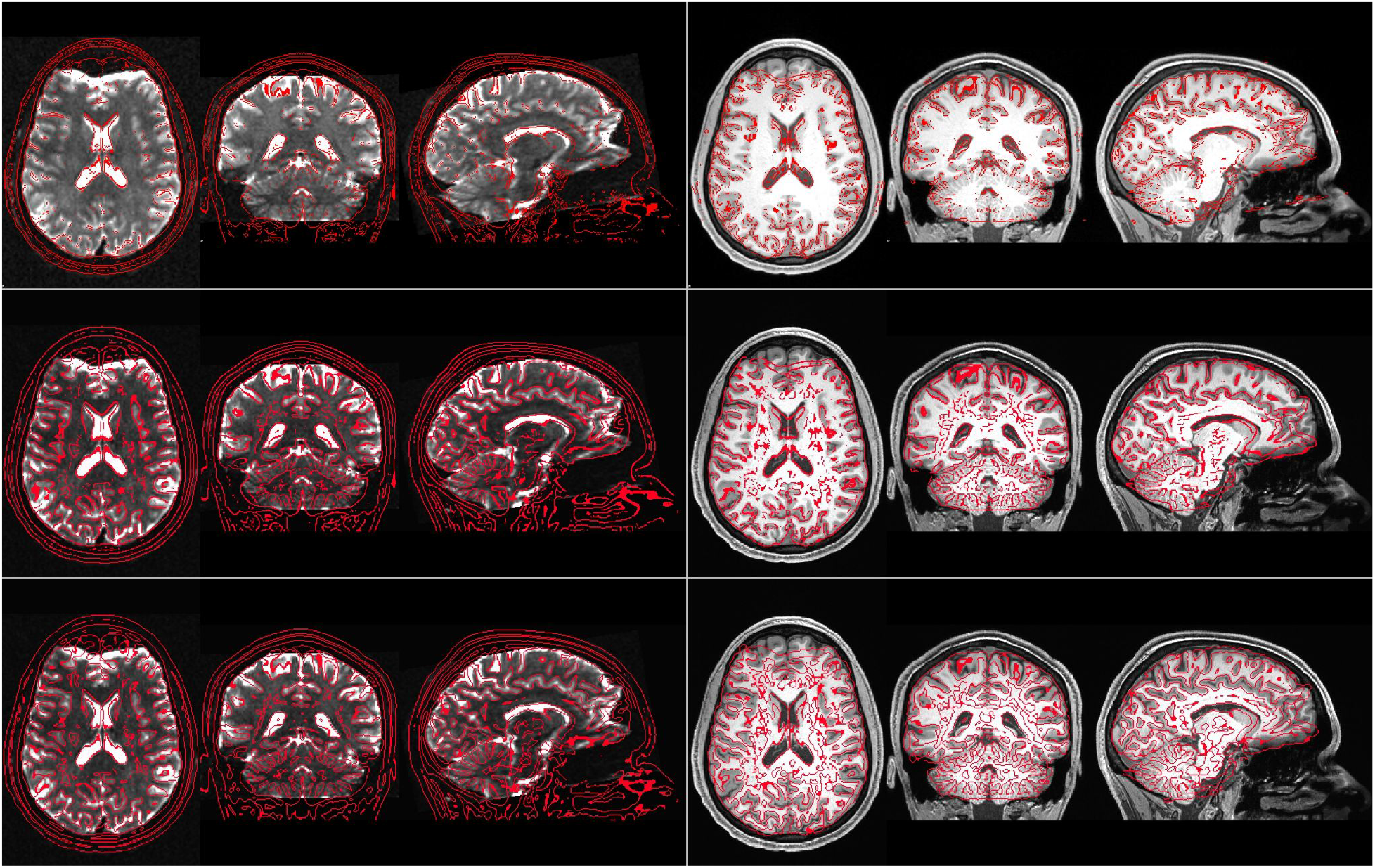
Example of rigid-body coregistration of a subject’s DWI (b=0 s/mm^2^) to a T1w volume using different edge approaches (from top row, downward): FSL’s slicer (first derivative-based method), AFNI’s 3dedge3 (Canny-based method) and AFNI’s 3dedgedog (new DoG-based method, with sigma_rad=1.4 and ratio_sigma = 1.4). See Figure 3’s caption for remaining details.

**Figure 5.**
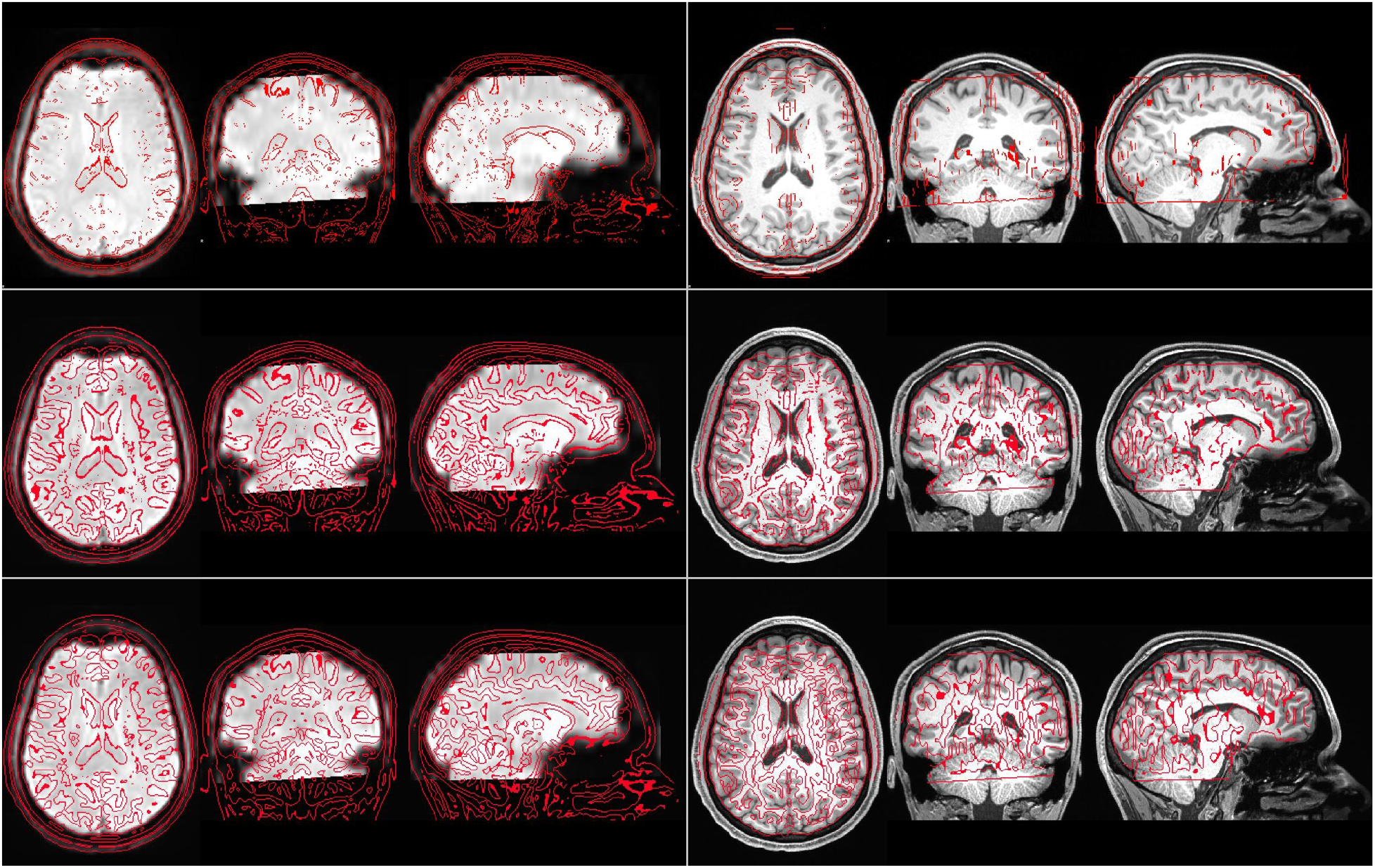
Example of rigid-body coregistration of a subject’s ASL volume to a T1w volume using different edge approaches (from top row, downward): FSL’s slicer (first derivative-based method), AFNI’s 3dedge3 (Canny-based method) and AFNI’s 3dedgedog (new DoG-based method, with sigma_rad=1.4 and ratio_sigma = 1.4). See Figure 3’s caption for remaining details.

In each image, the largest image contrasts occur at CSF, the skull and other facial feature boundaries, and each algorithm shows these clearly. However, the sulcal and gyral features, which are most important for judging meaningful brain feature alignment, appear differently across algorithms. Notably, the top row in each case shows a sparse representation of features, missing many edges and producing difficult to interpret maps.

The middle and bottom panels show remarkably similar representations, with most features clearly present in each and therefore beneficial for judging coregistration across the volume. This allows one to verify locations of good alignment---for example, most posterior sulci and gyri across modalities---as well as to identify those of poor alignment---such as the anterior regions where EPI distortion is present, as well as in the DWI’s ventricles. In general, the DoG method produces finer edges, allowing for more detailed comparisons of boundaries; for example, comparison T1w edges at the ventricle boundaries. For all methods, the edges derived from the EPI methods are noisier, as expected; the Canny method (middle panel) shows less noise within WM and clearly delineates major boundaries, but the DoG method appears to highlight more and finer features of the sulcal and gyral patterns.

### Additional edge estimation considerations

Both niimath and 3dedgedog provide the user with options to generate edges in a slicewise (2D) manner, which can be particularly useful and efficient for standard slicewise viewing of volumetric data. Figure 6 demonstrates this, using the sub-001’s anatomical dataset as input, using 3dedgedog. In the upper left panel, binary edges are estimated volumetrically using a standard sigma_rad=1.2 and ratio_sigma=1.2,. Note how some lines are particularly thick, representing out-of-plane structures that are tangential to the currently viewed slice. In the second row left panel, the “-only2D sag” option is used, showing cleaner lines within the slice; this facilitates views within the slice, and is particularly convenient for interactive displays and refreshing datasets (as the 2D estimation is faster to calculate). For static images, the 3D-derived edges may still be preferable.

**Figure 6.**
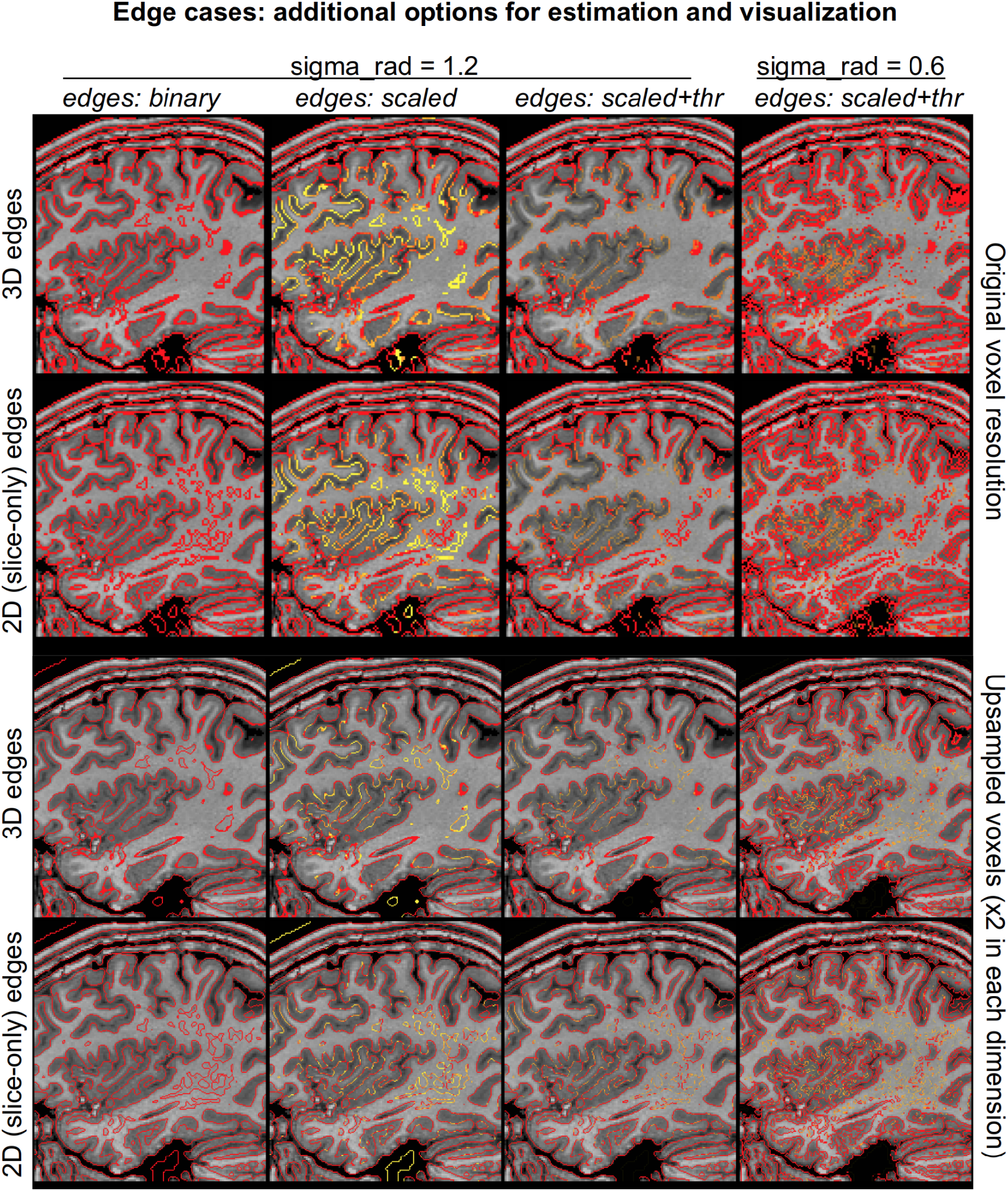
Additional considerations and features for edge estimation and visualization with 3dedgedog: 2D planar (slicewise) edges, boundary scaling and thresholding, and upsampling. The top-left panel shows standard, volumetric edges estimated for sub-001’s T1w dataset. The second row-left panel shows edges estimated only within the viewed sagittal plane: note that the thickest lines from the top panel, which appear when the slice is tangential to a structural boundary, do not appear, providing a clearer view of the given structures. The second column shows the edges when scaling (according to normalized local variation, red being highest) is applied, and in the third column this is translucently thresholded---thus, edges mostly likely to be “noise” are filtered. In the fourth column a smaller Gaussian is used to show more details, with both scaling and thresholding applied. The bottom two rows show the same types of edges, but first the input data has been upsampled by a factor of two in each dimension---the finer lines helpfully reduce image clutter, particularly when smaller sigma_rad is used.

In the second column of Figure 6, each boundary point is scaled by a measure of normalized local variation, ranging from 1 (yellow) to 100 (red). This information can be useful for removing or obscuring boundaries that are likely to be considered noise. Translucent thresholding of these scaled edges is applied in the AFNI viewer in the third column, leaving the main structural features primarily intact. In the fourth column, similarly scaled and thresholded edges are shown again, for the case of smaller sigma_rad=0.6. In this last column, both the gray-white and pial boundaries of the cortex are observable, while much of the “noise” of intra-WM variation is filtered out.

The bottom two rows of Figure 6 show edges created in the same manner as those in the top two rows, but with the difference that the anatomical dataset was first upsampled with 3dresample prior to edge estimation. The voxel edge length was halved in each dimension (so the output dataset has 8 times as many voxels), and nearest neighbor (NN) interpolation was used to avoid blurring the data. In this case, the edges can now be represented more finely, which is particularly important when viewing those estimated with sigma_rad=0.6 in the fourth column. In this case, the finer details can generally be appreciated more, and the image looks less cluttered. The computational cost is much higher, so this is less feasible for interactive updates, but still only takes about 10 seconds to estimate on a desktop.

## Discussion

We described and tested a simple and robust method for identifying edge boundaries on MRI scans. The approach relies on the Difference of Gaussian (DoG) algorithm. While some existing methods do produce useful edges (such as the Canny method), we suggest that the DoG method has some particularly useful properties for neuroimaging, particularly for registration assessment. First, it generates closed loops, even in noisy images, that help observers inspecting 2D slices to recognize the shape of 3D structures (and therefore their anatomical identity) more easily than disjointed lines. Additionally, the major adjustment parameter of the DoG is based on the spatial scale of the features of interest (which is typically known for a given MRI scan) rather than on the image’s relative intensity (which may vary across different regions of the brain and from scan-to-scan).

The spatial resolution tuning provides useful consistency across images that can have differing brightness inhomogeneities across scanners and modalities. This does mean, however, that the adjustment parameter must be adjusted/tuned prior to application in different species, for example, the monkey or rat brain, where the features would often be much lower than the 2mm scale used for humans. Depending on the spatial resolution and feature size, the blur sigmas could be adjusted. While we particularly focus on the known feature size of anatomical structures, it is worth noting that the spatial resolution of many sequences is adjusted to preserve a reasonable contrast to noise ratio. In other words, modalities with inherently low contrast to noise are often acquired with low spatial resolution (sampling a larger volume of hydrogen). Likewise, since signal increases with field strength, spatial resolution is often correlated with this attribute. This suggests an alternative (and cross-species) method for using the DoG spatial filter, which is based on the size of the voxels for each image; this is possible in some of the implementations coded here. Both approaches leverage the fact that the DoG uses space rather than intensity as the fundamental tuning parameter.

The DoG-based approach detailed in this paper is particularly well-suited to the construction of vivid edges that clearly illuminate the quality of inter-modality coregistration (e.g. alignment between T2*-weighted functional MRI scans and T1-weighted anatomical MRI scans acquired from the same). Integration of this approach in existing brain imaging preprocessing pipelines has the potential to improve early detection of erroneous coregistration that can be propagated to later stages. Moreover, the ability of users to tune the algorithm using a simple, easy to understand biologically calculable parameter (thickness of feature that needs to be highlighted), makes this technique flexible enough to be used by researchers interested in edge detection within systems where certain key properties are already known/consistent across samples.

While our approach is ideal for its proposed application, we would note that other visualization approaches may be more useful—or at least highly complementary—for later stages of image processing such as alignment to a template, where the primary goal is to warp (often nonlinearly) the shape of an individual’s brain to match a standard reference dataset. Standard templates often have known tissue probability maps (i.e. the probability of encountering of gray matter, white matter and cerebral spinal fluid at different locations is known), and the quality of coregistration can be improved by simultaneously segmenting the individual’s brain to these tissues (Ashburner and Friston, 2005). For template alignment, the use of a priori tissue maps can provide nice edge maps for a ground-truth solution. The Conn toolbox (Whitfield-Gabrieli and Nieto-Castanon, 2012) is one software program that benefits from this approach. Likewise, boundary-based registration (Greve & Fischl, 2009) methods directly estimate tissue boundaries and edges based on these estimates are ideal for evaluating the performance of this method. Clearly, some edge-based methods are tailored for specific applications. We merely suggest that DoG is a robust edge finding method for 3D images that requires little a priori expectations. It may be easily added to other visualization and coregistration strategies.

The proposed DoG-based edge-detection method is easy to implement in currently available neuroimaging pipelines. Smoothing data with a Gaussian kernel is already a common step for most neuroimaging pipelines as it diminishes high-frequency noise, reduces the effective number of statistical comparisons and makes it possible for researchers to interrogate their data using standard parametric statistical tests that assume gaussian distributions (Worsley et al., 1996). Therefore, as most neuroimaging pipelines include optimized Gaussian smoothing, the implementation of DoG is straightforward.

Poldrack and colleagues (2019) note that SPM/MATLAB, FSL, AFNI and Python are heavily used in neuroimaging, with each mentioned in more than 1000 publications per year. We have provided open source DoG solutions for users of each of these tools. This allows users to leverage this edge finding method using the tools they are familiar with. Specifically, the AFNI tool ‘3dedgedog’ (https://github.com/afni/afni), the niimath ‘dog’ functions (https://github.com/rordenlab/niimath), the FSL fslmath ‘-dog’ (https://fsl.fmrib.ox.ac.uk/fsl/fslwiki), the Matlab/SPM spmDoG script (https://github.com/neurolabusc/spmDoG), and the Python/nibabel PyDog script (https://github.com/neurolabusc/PyDog). Finally, we provide a web-assembly based implementation that can be used in any modern web capable device (phone, table, computer), embedded with our NiiVue visualization tool (https://niivue.github.io/niivue-niimath/). This final example includes a live demo that provides a zero-footprint ability to test this algorithm.

In summary, the DoG-based approach is a computationally efficient method to estimate edges within biomedical and neuroimaging datasets. The edges themselves have the useful property of boundedness, which is typically appropriate and useful for medical imaging applications. As explored above, the primary parameters for controlling the outputs are size-based, rather than brightness-based, which is also typically convenient for the variety of images obtained across the field. Furthermore, having the approximate ability to report edges “down to a certain scale length” is often a useful way of examining an image. The method is also flexible, with benefits of simple up-sampling and edge-scaling shown to be useful in further balancing the dual goals of seeing more details and not being distracted by small, noisy features. This methodology has immediate benefits for coregistration validation, and we believe it will have other beneficial applications in other domains with 3D voxel-based datasets such as confocal mircroscopy.

## Acknowledgments

This research and writing of the manuscript was funded by National Institute on Deafness and Other Communication Disorders Grants P50 DC014664 (CD, JF, RNN, TH, CR), and by the NIMH and NINDS Intramural Research Programs (ZICMH002888) of the NIH, HHS, USA (PAT, DRG).

## Author Contributions Statement

CR wrote the first draft to the manuscript and developed the niimath, Matlab and Python implementations and Figure 1. TH developed the FSL implementation. PT developed the AFNI implementation, evaluated the parameter space and generated figures 2-6. All authors made substantial identifying limitations, applications and niches for the algorithms and evaluating the results of the competing algorithms.

## Footnotes

1 https://afni.nimh.nih.gov/pub/dist/edu/data/CD.tgz

## References

Ashburner J, Friston KJ. (2005) Unified segmentation. Neuroimage. 26(3):839–51. doi: 10.1016/j.neuroimage.2005.02.018.

Bähnisch C, Stelldinger P, Köthe U (2009) Fast and accurate 3D edge detection for surface reconstruction, in Proc. DAGM Symp. Pattern Recogn. (Jena, Germany), 111–120. DOI: 10.1007/978-3-642-03798-6_12

Brett M, Leff AP, Rorden C, Ashburner J (2001) Spatial normalization of brain images with focal lesions using cost function masking. Neuroimage 14(2):486–500. doi: 10.1006/nimg.2001.0845.

Canny, J. (1986) A Computational Approach for Edge Detection,” IEEE Trans. Pattern Anal. Machine Intell., vol. 8, no. 6, 679–698.

Chen JS, Huertas A., Medioni G. (1987) Fast Convolution with Laplacian-of-Gaussian Masks/ IEEE Transactions on Pattern Analysis and Machine Intelligence, 9(4) 584–590. DOI: 10.1109/TPAMI.1987.4767946

Cox RW. (1996) AFNI: Software for analysis and visualization of functional magnetic resonance neuroimages. Computers and Biomedical Research, 29:162–173, 1996.

Deriche R. (1987) Optimal edge detection using recursive filtering. International Journal of Computer Vision, 1: 167–187.

Felzenszwalb PF, Huttenlocher DP (2012). Distance Transforms of Sampled Functions. Theory of Computing 8:415–428.

Fischl B, A M Dale1 (2000) Meassuring the thickness of the human cerebral cortex from magnetic resonance images. Proc Natl Acad Sci USA. 97(20):11050–5. doi: 10.1073/pnas.200033797.

Glen, D. R., Taylor, P. A., Buchsbaum, B. R., Cox, R. W., & Reynolds, R. C. (2020). Beware (Surprisingly Common) Left-Right Flips in Your MRI Data: An Efficient and Robust Method to Check MRI Dataset Consistency Using AFNI. Frontiers in neuroinformatics, 14, 18. https://doi.org/10.3389/fninf.2020.00018.

Gonzalez-Castillo, J., Duthie, K. N., Saad, Z. S., Chu, C., Bandettini, P. A., & Luh, W. M. (2013). Effects of image contrast on functional MRI image registration. NeuroImage, 67, 163–174. https://doi.org/10.1016/j.neuroimage.2012.10.076

Gonzalez R, Woods R (1992) Digital Image Processing, Addison-Wesley Publishing Company. SBN-10: 9780133356724

Greve DN, Fischl B. (2009) Accurate and robust brain image alignment using boundary-based registration. NeuroImage. 48(1): 63–72. https://doi.org/10.1016/j.neuroimage.2009.06.060.

Huertas A, Medioni G. (1986) Detection of Intensity Changes with Sub Pixel Accuracy Using Laplacian-Gaussian Masks. IEEE Transaction on Pattern Analysis and Machine Intelligence, 8, 651-664. DOI:10.1109/TPAMI.1986.4767838

Marr D, Hildreth E. (1980) Theory of Edge Detection. Proceedings of the Royal Society of London. Series B, Biological Sciences, 207 (1167): 187–217, doi:10.1098/rspb.1980.0020, PMID 6102765

Monga O, Deriche R, Malandain G, Cocquerez JP. (1991) Recursive filtering and edge tracking: two primary tools for 3-D edge detection. Image and Vision Computing 4:9, 203–214.

Poldrack RA, Gorgolewski K, Varoquaux G (2019) Computational and informatics advances for reproducible data analysis in neuroimaging, Annu. Rev. Biomed. Data Sci. 2:119–38

Prewitt JMS. (1970) Object Enhancement and Extraction, in Picture Processing and Psychopictorics, Lipkin, B.S., and Rosenfeld, A. (eds.), Academic Press, New York.

Raslau F, Lin LY, Andersen A, Powell DK, Smith CD, Escott E. (2018) Peeking into the Black Box of coregistration in Clinical fMRI: Which coregistration Methods Are Used and How Well Do They Perform? American Journal of Neuroradiology. 39(12): 2332–2339.doi: 10.3174/ajnr.A5846

Roberts L. (1965) Machine Perception of 3-D Solids, Optical and Electro-optical Information Processing, MIT Press 1965.

Saad ZS, Glen DR, Chen G, Beauchamp MS, Desai R, Cox RW (2009). A new method for improving functional-to-structural MRI alignment using local Pearson correlation. Neuroimage 44 (3), 839–848

Smith SM, Jenkinson M, Woolrich MW, Beckmann CF, Behrens TE, Johansen-Berg H, Bannister PR, De Luca M, Drobnjak I, Flitney DE, Niazy RK, Saunders J, Vickers J, Zhang Y, De Stefano N, Brady JM, Matthews PM (2004). Advances in functional and structural MR image analysis and implementation as FSL. Neuroimage 23:S208–S219.

Sobel IE. (1970) Camera Models and Machine Perception, Ph.D. dissertation, Stanford University, Palo Alto, CA.

Spontón H, Cardelino J. (2015) A Review of Classic Edge Detectors. Image Processing On Line, 5(2015), 90–123. https://doi.org/10.5201/ipol.2015.35

Whitfield-Gabrieli S, Nieto-Castanon A. (2012) Conn: a functional connectivity toolbox for correlated and anticorrelated brain networks. Brain Connect. 2(3):125–41. doi: 10.1089/brain.2012.0073.

Wilson HR, Giese SC. (1977). Threshold visibility of frequency gradient patterns. Vision Research, 17(10), 1177–1190. https://doi.org/10.1016/0042-6989(77)90152-3

Worsley KJ, Marrett S, Neelin P, Vandal AC, Friston KJ, Evans AC. (1996) A unified statistical approach for determining significant signals in images of cerebral activation Human Brain Mapping, 4:58–73.

